# Interactive effects of shifting body size and feeding adaptation drive interaction strengths of protist predators under warming

**DOI:** 10.1101/101675

**Authors:** K. E. Fussmann, B. Rosenbaum, U. Brose, B.C. Rall

**Affiliations:** J.F. Blumenbach Institute of Zoology and Anthropology, University of Göttingen, Berliner Str. 28, 37073 Göttingen, Germany; German Centre for Integrative Biodiversity Research (iDiv), Halle-Jena-Leipzig, Deutscher Platz 5e, 04103 Leipzig, Germany; Institute of Ecology, Friedrich Schiller University Jena, Dornburger-Str. 159, 07743 Jena, Germany

**Keywords:** climate change, functional response, body size, temperature adaptation, activation energies, microcosm experiments, predator-prey, interaction strength, Bayesian statistics, Paper type: Primary Research

## Abstract

Global change is heating up ecosystems fuelling biodiversity loss and species extinctions. High-trophic-level predators are especially prone to extinction due to an energetic mismatch between increasing feeding rates and metabolism with warming. Different adaptation mechanisms such as decreasing body size to reduce energy requirements (morphological response) as well as direct effects of adaptation to feeding parameters (physiological response) have been proposed to overcome this problem. Here, we use protist-bacteria microcosm experiments to show how those adaptations may have the potential to buffer the impact of warming on predator-prey interactions. After adapting the ciliate predator *Tetrahymena pyriformis* to three different temperatures (15°C, 20°C and 25°C) for approximately 20 generations we conducted functional response experiments on bacterial prey along an experimental temperature gradient (15°C, 20°C and 25°C). We found an increase of maximum feeding rates and half-saturation densities with rising experimental temperatures. Adaptation temperature had on average slightly negative effects on maximum feeding rates, but maximum feeding rates increased more strongly with rising experimental temperature in warm adapted predators than in cold adapted predators. There was no effect of adaptation temperature on half-saturation densities characterising foraging efficiency. Besides the mixed response in functional response parameters, predators also adapted by decreasing body size. As smaller predators need less energy to fulfil their energetic demands, maximum feeding rates relative to the energetic demands increased slightly with increased adaptation temperature. Accordingly, predators adapted to 25°C showed the highest feeding rates at 25°C experimental temperature, while predators adapted to 15°C showed the highest maximum feeding rate at 15°C. Therefore, adaptation to different temperatures potentially avoids an energetic mismatch with warming. Especially a shift in body size with warming additionally to an adaptation of physiological parameters potentially helps to maintain a positive energy balance and prevent predator extinction with rising temperatures.

## Introduction

Global change has a negative impact on biodiversity, up to a point where scientists consider the world to be on the verge of the sixth wave of mass extinction (Wake & Vredenburg, 2008; Pereira *et al*., 2010; Barnosky *et al*., 2011). Changing temperatures are one major driver of global change and are expected to have a global impact (MEA, 2005). Climate reports predict a minimum increase of 1.5°C in surface temperature by the end of the century and it is deemed extremely likely that anthropogenic causes (Cook *et al*., 2013) have led to the warmest 30 year period of the last 1,400 years (IPCC, 2014). Temperature directly affects development, survival, range and abundance of species (Bale *et al*., 2002) and has a strong effect on species interactions (Montoya & Raffaelli, 2010) as well as on the structure and dynamics of species communities (Brose *et al*., 2012). Further, increasing temperatures in aquatic as well as terrestrial ecosystems have been linked to vast biodiversity losses during extinction waves in earlier earth periods (Gómez *et al*., 2008; Mayhew *et al*., 2008; Joachimski *et al*., 2012). Despite this negative impact of high temperatures on taxonomic richness, previous periods of warming have also been associated with high speciation rates since increasing temperatures can trigger rapid evolution (Gillooly *et al*., 2005; Geerts *et al*., 2015) and create niche openings by eliminating species previously occupying a certain habitat or resource (Mayhew *et al*., 2008). This leads to the question whether adaptation and evolution pose a feasible escape from warming induced extinction.

Species’ extinctions and therewith biodiversity strongly depend on the stability of the ecosystems they are embedded in (May, 1972; McCann, 2000). Stability furthermore depends on the interaction strengths between species. High interaction strengths decreases the population stability leading to extinction caused by high population cycles, and too low an interaction strength may lead to extinction of predators due to starvation (Rall *et al*., 2010).

The functional response is one of the oldest and most established tools to quantify the strength of these interactions and describe species-species feeding interactions in ecology (Holling, 1959; Jeschke *et al*., 2002). In this framework, the feeding rate, *F*, of a predator depends on the density of its resource. The functional response, as described by Real (1977) includes a non-linear feeding rate, which determines the maximum feeding, *f*, when prey is abundant. At lower prey densities the functional response curve is characterised by the predator’s foraging efficiency. Mathematically this is described by half-saturation density, *ƞ*, the prey population density at which half of the maximum feeding rate is reached. (Figure 1). These parameters can be used to evaluate interspecies interaction strength which have been a main predictor of ecosystem stability (Berlow *et al*., 2009).

**Figure 1:**
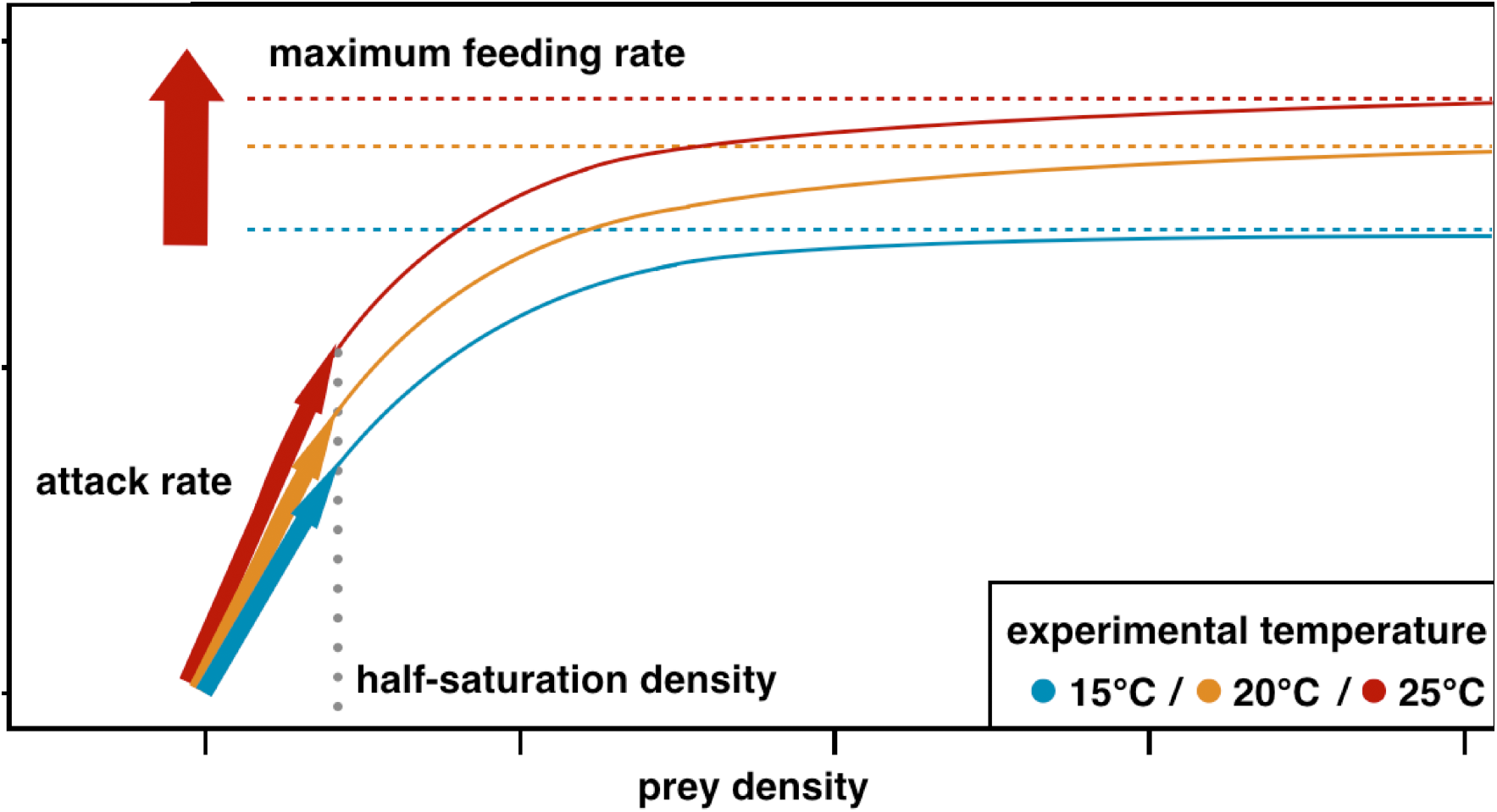
Expected trends for changes in maximum feeding rate (dashed lines) and half-saturation density (dotted line)with increasing temperatures based on previous studies (Rall etal., 2012; Fussmann etal., 2014). Maximum feeding reates are likely to increase with experimental temperature, which half saturation densities have a variable scaling relationship being on average neutral. Maximum feeding rates of predators adapted to higher temperatures are expected to increase to cope with increasingmetabolic demands with the highest maximum feedingg rates at high temperatures and vice versa for coldadapted predators. Half saturation desities are expected to not be influenced by temperature adaption resulting in an increase of attack rates(arrows) at low prey densities in warm adapted predators to facillate higher maximum feeding rates at high prey densities.

To investigate the effects of warming on interaction strength, previous studies have used the principles of the Metabolic Theory of Ecology (MTE) (Gillooly *et al*., 2001; Brown *et al*., 2004) which is quantified as activation energies measured in electron Volt [eV] according to the Arrhenius equation (Arrhenius, 1889). The Arrhenius equation, originally used to describe chemical reactions and enzyme kinetics, has become a mechanistic model for biological rates in ectotherm organisms (Gillooly *et al*., 2001; Brown *et al*., 2004; Savage *et al*., 2004). The MTE argues that all biological rates as well as higher order patterns such as density distributions scale with temperature. Therefore, the parameters of the functional response, determining interaction strength should follow the same principles (Vasseur & McCann, 2005; Fussmann *et al*., 2014): maximum feeding rates are often assumed to scale with temperature in the same manner as metabolic demands with an activation energy ranging from 0.6 to 0.7 eV across different taxa (Vasseur & McCann, 2005). This is corroborated by empirical data of ciliates, flagellates and other microfauna showing even higher activation energies for maximum feeding rates of 0.772 eV (Hansen *et al*., 1997; Vasseur & McCann, 2005). However, a broader analysis of predators from different ecosystems revealed that maximum feeding, *f*, scales with an activation energy of roughly 0.3 to 0.4 eV (Rall *et al*., 2012; Fussmann *et al*., 2014). This leads to a mismatch where, under warming, metabolism increases faster than maximum feeding. As a result, predators cannot meet their metabolic demands and run the risk of starvation even if they are surrounded by prey (Vucic-Pestic *et al*., 2011; Fussmann *et al*., 2014). The half-saturation density, *η*, can be influenced by a variety of parameters such as encounter rate, mobility of prey and predator, and search efficiency. Most significantly, mobility of prey and predator and therefore encounter rates and search efficiencies are influenced by warming (Sentis *et al*., 2012; Dell *et al*., 2014). Since the reaction of prey as well as predator to changing temperatures can be highly variable in both, general tendency and intensity, this results in a high variability of activation energies for half-saturation density, ranging from positive to negative relationships with warming, being on average neutral (Fussmann *et al*., 2014). Constant half-saturation densities can be mechanistically explained by a simultaneous increase of feeding at low densities (the “rate of successful attacks” (Holling, 1959), often referred to as attack rate, capture rate or maximum clearance rate) and maximum feeding rate with increasing temperatures. At high temperatures, natural systems show lower prey population densities due to reduced resource availability (Brown *et al*., 2004; Meehan, 2006; Fussmann *et al*., 2014). If predator abundances are low in natural systems and predators are not able to increase their foraging efficiency under those conditions (i.e. decrease of half-saturation densities) feeding rates eventually decrease. These mismatches can lead to the loss of higher trophic levels due to starvation (Binzer *et al*., 2012; Fussmann *et al*., 2014) and decreases in biodiversity due to warming (Binzer *et al*., 2016).

However, other mechanisms, for example adaptation to higher temperatures, may be able to counteract starvation due to energetic mismatch (Angilletta Jr., 2009; Chevin *et al*., 2010; Somero, 2010). In studies where adaptation may have buffered the physiological impacts of warming, temperature had hardly any effect on the overall fitness of a population (Chown *et al*., 2010). Adaptation in predator-prey systems is often studied from a prey′s perspective (McPeek *et al*., 1996; Yoshida *et al*., 2003; Abrams & Walters, 2010) but rarely from a predator's (Sentis *et al*., 2015), despite them being most affected by temperature changes (Rall *et al*., 2010; Binzer *et al*., 2012; Fussmann *et al*., 2014). Given that predator energy efficiency is a major determinant of population stability (Vasseur & McCann, 2005; Rall *et al*., 2010), an adaptation of either metabolism or functional response parameters or both could be crucial. Temperature adaptation, however, is often investigated on short time scales (Sentis *et al*., 2015) leading to concerns that the time frame of temperature changes exceeds adaptation rates (Quintero & Wiens, 2013). Generally, short-term studies tend to underestimate a species’ capability of adapting to climate change (Leuzinger *et al*., 2011). In a short-term study focussing on acclimation within one generation, the physiological temperature effect on feeding rates proved crucial since metabolic rates and body size were less affected by acclimation temperature (Sentis *et al*., 2015). However, metabolism and functional response parameters are not only influenced by temperature but also by body size (Vucic-Pestic *et al*., 2011; Ott *et al*., 2012; Rall *et al*., 2012; Kalinkat *et al*., 2013). However, body size itself is influenced by temperature (Atkinson *et al*., 2003; Figure 2). Globally, species in warmer regions tend to have smaller average body sizes than species in colder ecosystems (Bergmann, 1847), this trend was also documented in warming studies investigating different size spectra of local freshwater communities (Daufresne *et al*., 2009; Yvon-Durocher *et al*., 2011). Further, body size has been shown to have a strong effect on interaction strengths through allometric scaling (Brose, 2010). Smaller body sizes require less energy to maintain metabolism and population growth (Brown *et al*., 2004) leading to reduced maximum feeding rates while not affecting the half-saturation densities (Hansen *et al*., 1997). The half-saturation density (*η* = 1/*(T*_*h*_*a))* can be calculated as the inverse of the product of handling time (*T*_*h*_*=1/f*) and attack rate (*a = f/η*). Consequently, if maximum feeding rates and attack rates scale similarly with body size, the effect on half-saturation density is equalled out (*η = f/a)* (Rall *et al*., 2012). As a result of constant half-saturation density, maximum feeding rates are constant across the entire prey density gradient (Figure 1).

**Figure 2:**
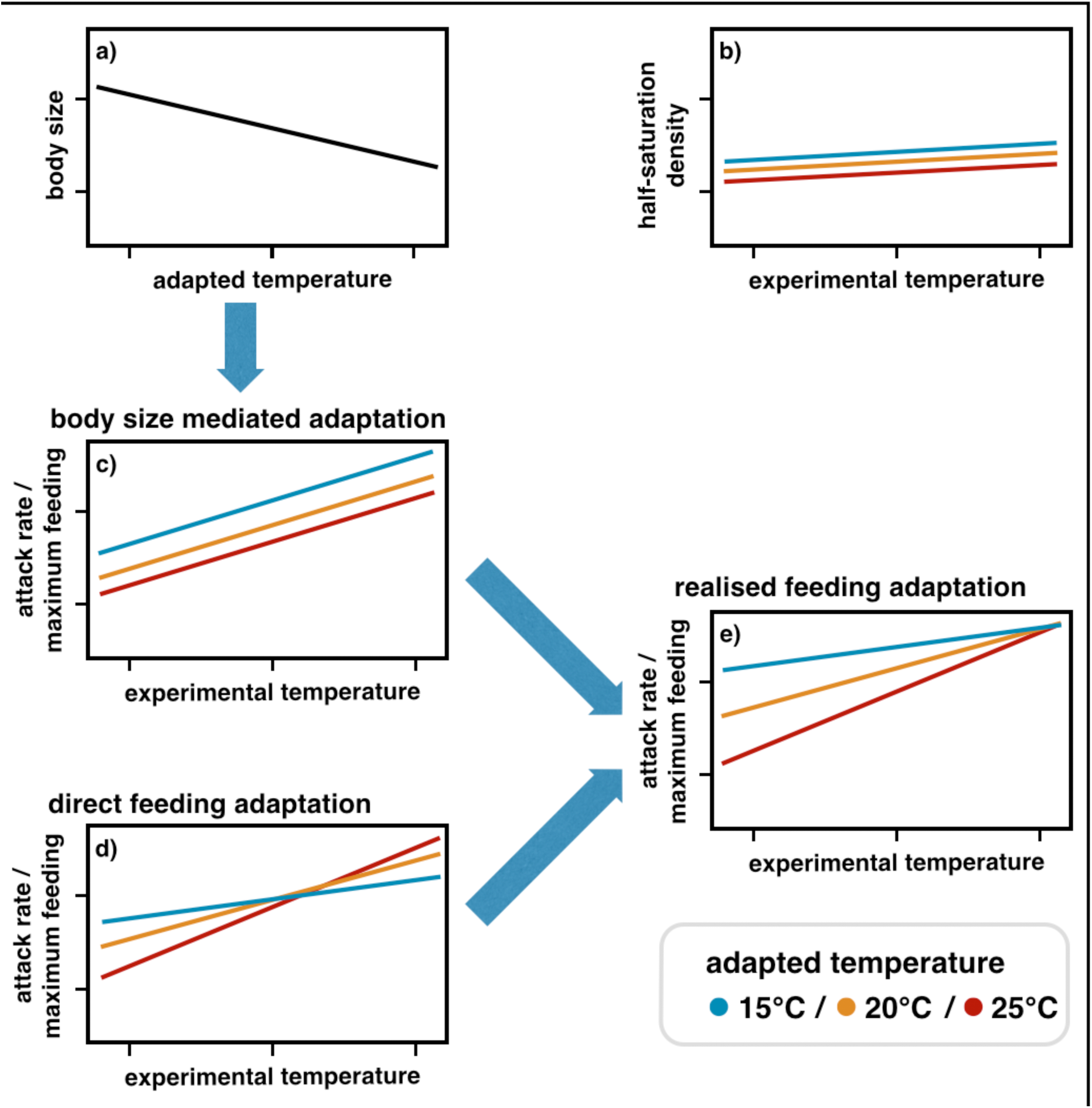
**a) Predator body size** (cell size) decreases with increasing adaptation temperature. **b) Half-saturation densities** are not expected to change with experimental or adaptation temperature. **c) Body size mediated adaptation**: Decreasing predator body size with increasing adaptation temperatures generally reduces maximum feeding rates in warm adapted predators with an assumed scaling exponent of 0.75 (Brown *et al*., 2004). **d) Direct feeding adaptation**: Maximum feeding rates generally increase with rising experimental temperatures. Adaptation to temperature leads to a direct feeding adaptation of maximum feeding rates. Predators adapted to 15°C are expected to show the overall highest maximum feeding at 15°C experimental temperature, while predators adapted to 25°C should have the overall highest maximum feeding rates at 25°C. **e) Realised feeding adaptation:** Realised feeding adaptation shows the interactive effects of body size mediated adaptation and direct feeding adaptation on maximum feeding rates.

Here, we explored how interactive effects of direct temperature adaptation of feeding rates and indirect effects on feeding rates through temperature induced changes in body size influence functional response parameters. We designed a microcosm experiment with short generation times (Callahan *et al*., 2008) to understand how adaptation to different temperatures over 20 generations influences feeding behaviour. We investigated whether adaptation to temperature enables predator populations to avoid extinction caused by crossing the threshold where metabolic demands overtake the energy intake through feeding. (1) We expect body sizes of warm adapted *Tetrahymena* to decrease within 20 generations compared to predators adapted to colder temperatures (Figure 2a). (2) Half-saturation densities should not be affected by increasing experimental temperature. If *Tetrahymena* adapts both, attack rates and maximum feeding rates simultaneously, we expect no change in half-saturation density with adaptation temperature (Figure 2b). (3) The change in body size, (cell size) will cause a decrease in maximum feeding rates and attack rates in warm adapted predators. Body size, however, will not affect the temperature dependency of these rates or change the activation energies (Figure 2c – body size mediated feeding adaptation). (4) We assume thatthe direct physiological adaptation of maximum feeding rates and attack rates leads to a change of activation energies: warm adapted predators should show the steepest increase (highest activation energy) as they should be well adapted to higher temperatures. Predators adapted to lower temperatures should have the highest feeding rates at cold temperatures but will not or just marginally be able to increase feeding with increasing temperature (Figure 2d – direct feeding adaptation). These different scalings will result in a statistically significant interaction. (5) If both mechanisms occur simultaneously, maximum feeding rates and attack rates should be lowest for warm adapted predators and increase with decreasing adaptation temperature while keeping the interactive direct effect of adaptation (Figure 2e – realised feeding adaptation).

## Methods

### Laboratory Cultures

We chose a model predator-prey system with the non-toxic bacterium *Pseudomonas fluorescens* CHA19 (Zuber *et al*., 2003; Jousset *et al*., 2009) as prey and the ubiquitous, predatory ciliated protozoan *Tetrahymena pyriformes* CCAP 1630/1W (CCAP Culture Collection of Algae and Protozoa, SAMS Limited, Scottish Marine Institute, Scotland, United Kingdom).

*Pseudomonas fluorescens* CHA19 was marked with GFP using a Mini-TN7 transposon I (Lambertsen *et al*., 2004). After molecular cloning, one colony of the *Pseudomonas* strain was deep frozen at -80 °C in a 25 % glycerol solution. For every experiment a small sample was defrosted and incubated on LB-Agar containing 8 µg/l gentamycin before single colonies were incubated at room temperature in selective LB-medium over night. *Tetrahymena* was grown in 2% proteone peptose medium at 20°C. At the start point of the adaptation experiment, the culture of *Tetrahymena* was divided equally into 9 cultures, 3 cultures were henceforth kept at 15°C, three cultures were kept at 20°C and three cultures were kept 25°C. A temperature range between 15°C and 25°C is realistic for temperate aquatic systems in absence and presence of an extreme temperature event (Seifert *et al*., 2015). For all adaptation temperatures, exponential growth rates of *Tetrahymena pyriformis* were measured to estimate the timeframes until approximately 20 generations were reached and functional response experiments were conducted (Supporting Information Figure 1). For predators kept at 15°C adaptation temperature, this was approximately 18 days, while for warmer adapted predators this time span was approximately 13 and 12 days for 20°C and 25°C adaptation temperature, respectively. 20 generations is consistent with other studies ranging from only one generation (Sentis *et al*., 2015) to 10 and 100 generations (Padfield *et al*., 2015). To reduce the traces of medium prior to the functional response measurements bacteria were centrifuged (13.000 rpm x 1 min) and re-suspended three times in sugarless Ornston and Stannier (OS) medium (Ornston, 1966) diluted with ddH2O 1:10. Bacterial counts were measured using an Accuri C6 flow cytometer (BD Biosciences) on slow with an FSC-H of 8000 and a SSC-H of 2000. Ciliates were harvested by centrifugation at 300 rpm for 7 min at 0°C and re-suspended in OS medium three times. Prior to functional response experiments the predators were starved for 12 hours at their respective adaptation temperatures. The number of ciliates and their body sizes were measured with a Beckman Coulter Counter Multisizer 4 with a 100µl aperture on slow fluidics speed (Beckman Coulter, Inc.).

### Functional Response Experiments

Functional response experiments were conducted in 96-well plates. One column contained only the ciliated predator *Tetrahymena*, each of the remaining 11 columns contained a different bacterial prey density, with six rows as identical replicates and two rows as bacterial controls. Each well contained 200µl of sugar-free OS 1:10 media, prey densities ranged from 34778 bacteria µl^-1^ to 1189416 bacteria µl^-1^, while predator abundances were kept constant at 100 predators µl^-1^. We used a fully factorial design for the functional response experiments, conducting experiments at the full experimental temperature range of 15°C, 20°C and 25°C with all three cultures of all adaptation temperatures after approximately 20 generations (Supporting Information Figure 1). Fluorescence intensities of bacteria were measured at two time points, after four hours into the experiment to avoid transient dynamics and at the end of the experiment, three hours thereafter, in an Infinite M200 plate reader (Tecan, Männedorf, Switzerland). After orbital shaking for 10 seconds, each well was measured with an excitation wavelength of 485 nm and an emission wavelength of 520 nm reading 15 flashes with a manual gain of 100. To standardise a reliable value for bacterial abundance from the GFP signal measured in the plate reader, comparative measurements were taken using a plate reader and flow cytometer with an FSC-H of 8000 and SSC-H of 2000 and slow fluidics speed.

### Calculation of bacterial densities

We assessed bacterial fluorescence data by using a regression tree (tree-function, Ripley, 2016) classifying count-fluorescens relationships (Supporting Information, Figure 1).To estimate the bacterial density we first fitted the ln-transformed fluorescence signal measured in the plate reader against the ln-transformed number of cells measured with the flow cytometer (independent variable). All fluorescence values below a ln(GFP) of 6.03068 and a ln(count) of 9.071045 and above a ln(GFP) of 9.37609 and a ln(count) of 13.15602 were excluded from further analysis since we could not guarantee the proportionality between cell count and fluorescent signal beyond these counts. We then calculated bacterial abundance by predicting a linear model with GFP signal and experimental temperature as independent variables. To account for background signals, all experimental data was blanked against OS 1:10 ddH_2_O experimental media and treatments containing only the predator *Tetrahymena pyriformis*. This resulted in 141 control treatments containing only bacterial prey, and 306 functional response experiments that were used for further analysis (Supporting Information Table 1).

**Table 1:** Models for including experimental and adaptation temperature. All models include ln-linear terms of experimental temperature in the scaling of *f*, *ƞ*, *r*, *K* (equations 4a-d).

| Model | Equations | Terms for influence of adaptation temperature |
| --- | --- | --- |
| 1 | (4a), (4b) | none |
| 2 | (5a), (5b) | ln-linear in $f$ and $\eta$ |
| 3 | (6a), (5b) | interaction with experimental temperature in $f$, ln-linear in $\eta$ |
| 4 | (5a), (6b) | ln-linear in $f$, interaction with experimental temperature in $\eta$ |
| 5 | (6a), (6b) | interaction with experimental temperature in $f$ and $\eta$ |

### Functional response

The functional response describing the non-linear feeding rate, *F*, is defined as (Real (1977):

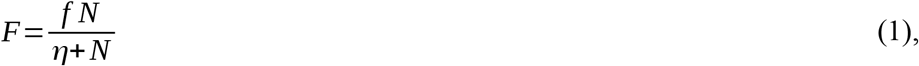

where *N* is the prey density, *f* is the maximum feeding rate and *η* is the half-saturation density. In our experiment, additional to a constant decline of prey through time due to feeding, natural growth and mortality of the bacterial prey occurred in control experiments. We therefore decided to incorporate a Gompertz growth for microbiological systems (Gompertz, 1825; Paine *et al*., 2012):

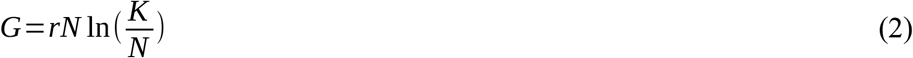

where *r* is the intrinsic growth rate of bacteria and *K* is the carrying capacity of bacteria. A model accounting for changes in prey abundance over time due to feeding as well as natural prey growth or death is expressed in the following ordinary differential equation (ODE):

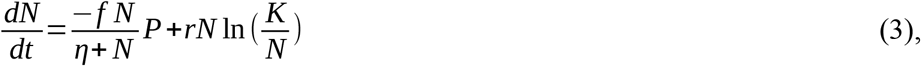

where the change in prey abundance over time *t* is characterised by the functional response model; *P* is the predator density. To account for changes of the parameters with experimental temperature we calculated Arrhenius temperatures and activation energies *E*_*f*_,
_*η*_:

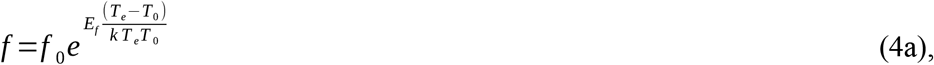

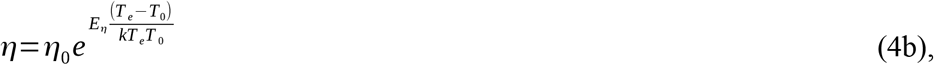

where *f*_0_ and *η*_0_ are normalisation constants, *T*_*e*_ [K] is the absolute experimental temperature, *T*_0_ [K] is the normalisation temperature and *k* [eV K^-1^] is the Boltzmann's constant yielding the well known Arrhenius equation (Arrhenius, 1889; Gillooly *et al*., 2001).

Additionally, growth and carrying capacity also scale with temperature (Gillooly *et al*., 2001; Savage *et al*., 2004):

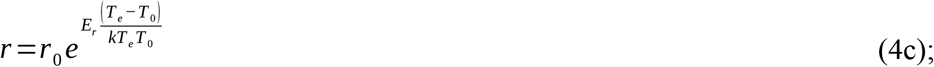

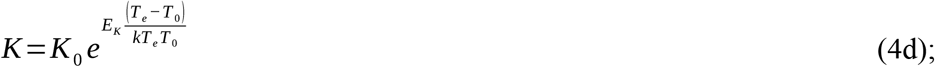

with *r*_0_ and *K*_0_ being normalisation constants, and *E*_*r*_ and *E*_*K*_ being the experimental activation energies. To investigate the effects of adaptation, we extended the Arrhenius equation using a term describing the dependency of the maximum feeding rate, *f*, and of the half-saturation density, *η*, on the temperature the predator was adapted to *T*_*a*_:

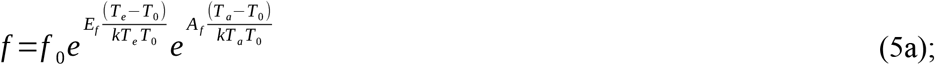

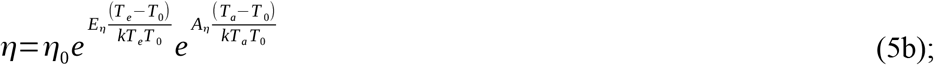

where *A*_*f*_ and *A*_*η*_ are the activation energies for temperature adaptation. Both, maximum feeding and half-saturation density may interactively react to both, experimental and adaptation temperature (i.e. E_*f*_,_*η*_ is different for different *T*_*a*_). We therefore introduced an interaction term, *I*_*f*_,_*η*_, into equation 5a, b (i.e. statistical interaction term), yielding:

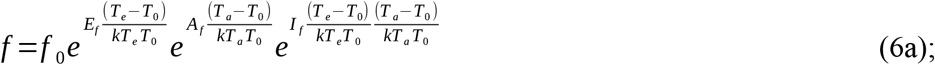

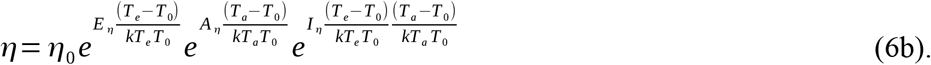

Further, we calculated realised activation energies for maximum feeding rates

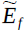
 of the experimental temperature for each adaptation temperature:

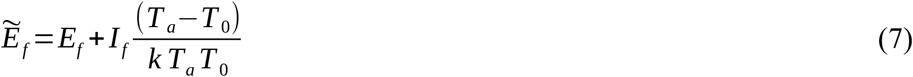

Maximum feeding rates, *f*, scale not only with temperature but also with body size with a power-law exponent of 0.75 according to the MTE (Yodzis & Innes, 1992; Brown *et al*., 2004). Half-saturation densities, *η*, can be defined as the quotient of maximum feeding rate and attack rate (*η* = *f*/*a*, Koen-Alonso, 2007), where both parameters share the same power-law exponent of 0.75 (Brown *et al*., 2004) and do not scale with body size (Yodzis & Innes, 1992; Hansen *et al*., 1997).

The body size dependent functional response can therefore be described with a ¾ power law scaling of the maximum feeding rate, *f*, with body size, *m*:

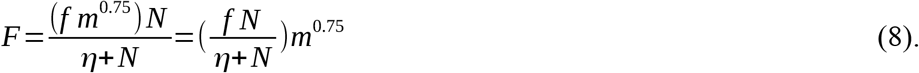

To demonstrate the effect of direct feeding adaptation (Figure 2d – direct feeding adaptation), we corrected our fitted results based equation on 5a (see below and in the Supporting Information a description of the fitting methods) by dividing feeding rates by the metabolic body size of the predator (Schmitz & Price, 2011; Schneider *et al*., 2012):

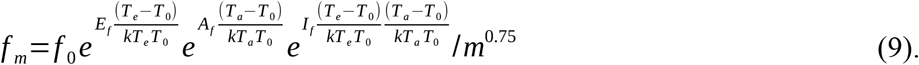

Mean ciliate body size [µm^3^], adapted to the respective temperatures at the time of experiment after an adaptation period of approximately 20 generations, was measured in the Beckmann Coulter Counter. Note that this calculation was done after fitting the functional response model to the data. This method to correct for body size differences in temperature dependent functional response parameters was already successfully applied in prior studies (Sentis *et al*., 2012, 2014).

### Fitting algorithm

We used Bayesian methods for parameter estimation (equation 3 including scaling relationships for *r*, *K*, *f* and *η*). Data of prey densities after 4 hours *N*(*t*_4_) were used as initial values for the numerical solution of the ordinary differential equations (ODE) and data of densities after 7 hours *N*(*t*_*7*_) were modelled using ln-normally distributed errors. Model parameters for control treatments and treatments with predators present were estimated within the same model. Samples from the posterior distribution of the parameters given the data were drawn using Hamiltonian Monte Carlo sampling in Stan, accessed via the RStan package (Stan Development Team, 2016). The Stan software comes with a built in ODE-solver, making it suitable for fitting ODE-based functional responses (equation 3). We used normally distributed uninformative priors with zero means and standard deviations of 100,000 for *K*_*0*_ and *ƞ*_*0*_, standard deviations of 100 for all other model parameters and a uniform distribution on the interval between 0 and 100 as a prior for the model's standard error. The parameters *r*_*0*_, *K*_*0*_, *f*_*0*_, and *ƞ*_*0*_ were provided with a lower boundary of zero. We ran 5 Markov chains in parallel with an adaptation phase of 1,000 iterations and 20,000 sampling iterations each, summing up to 100,000 samples of the posterior distribution. Visual inspection of the trace plots and density plots showed a good mixture of the chains. Values of

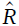
 sufficiently close to 1 and an adequate effective sample mass *n*_*eff*_ further verified *R* convergence (Supporting Information, Table 3). We tested different models for including adaptation temperature in the scaling relationships of *f* and *ƞ* (Table 1). For model comparison we used the Watanabe-Akaike information criterion (WAIC), which can be computed from the log-likelihood values of the posterior samples by the loo package (Vehtari & Gelman, 2016). We will report results only for model 3, which performed best in the model selection (Supporting Information, Table 2). Model 3 includes an interaction term of experimental and adaptation temperature in the scaling of maximum feeding rate *f*, but not in the scaling of half-saturation density *ƞ.* The fits of the full ODE together with the measured data points as well as functional response plots can be found in the Supporting Information (Supporting Information Figure 3 and 4). See Supporting Information also for full summary statistics, density plots and model code.

**Figure 3:**
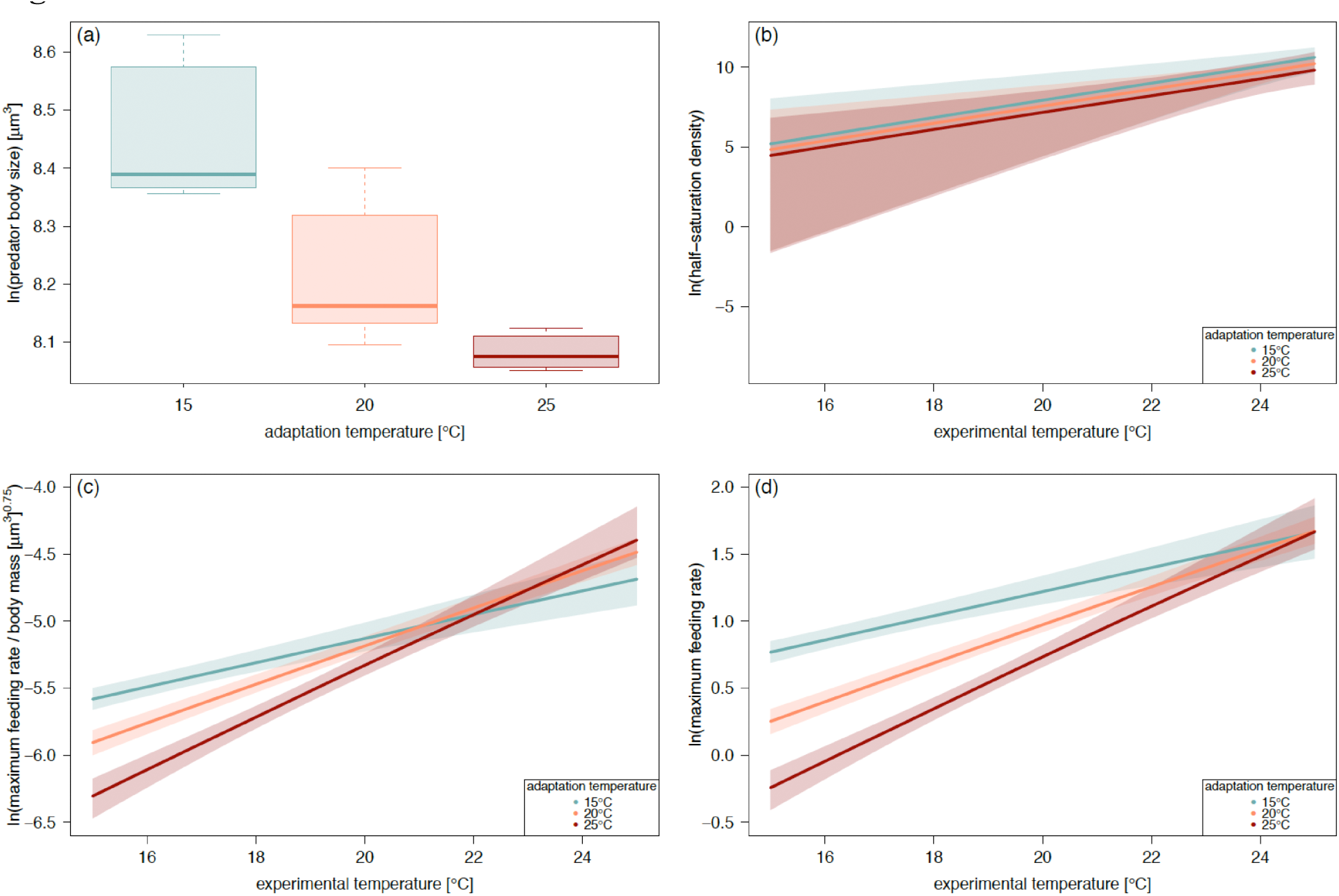
**a) Body sizes** of Tetrahymena pyriformis adapted to 15oc, 20oc,250c in um3 measured in the Beckmann coulter counter decreased with adaption temperatures. **b)Half-saturation densities** for *tetrahymena pyriformis* adapted to 15oc(blue0, 20oc(orange and 25oc(red))increased with experimental temperatures. There was no significant difference for half-saturation density between predators adapted to different temperatures along the gradient of experimental temperatures **c)Metabolic body size accounted maximum feeding rates**(f/body size0.75[um3])for *tetyrahymena pyriformis* adapted to15oc, 20oc, 250c along an experimental temperatures while predators adapted to 15oc and 20oc showed the highest maximum feeding rates at their adaption temperatures, respectively. **d)Maximum feeding rates** for *Tetrahymena pyriformis* adapted to 15oc, 20oc, 250c increased with experimental temperatures. While maximum feeding rates slightly decreased with adaptation temperatures, predators adapted to 25oc over 20 generations showed the strongest increase in maximum feeding with experimental temperature due to a positive interaction effect of experimental and adaptation temperatures. Solid line represent median values shaded areas indicate 95% credibility intervals.

**Table 2:** Mean values of the distribution and their standard deviation for normalisation constants of maximum feeding rate *f*_*0*_ and half-saturation density *ƞ*_*0*_ and their activation energy main effects of experimental temperature *E*_*f*_, *E*_*ƞ*_, of adaptive temperature *A*_*f*_, *A*_*ƞ*_, and the interaction term for maximum feeding rate *I*_*f*_. The range between 2.5% and 97.5% of the distribution give the 95% credible intervals. For full summary statistic, please see Supporting Information Table 3.

|  | mean | sd | 2.5% | 50% | 97.5% |
| --- | --- | --- | --- | --- | --- |
| $f_0$ | 2.654 | 0.085 | 2.505 | 2.647 | 2.841 |
| $\eta_0$ | 2378.247 | 1932.152 | 83.463 | 1930.298 | 7115.469 |
| $E_f$ | 1.054 | 0.056 | 0.950 | 1.052 | 1.168 |
| $E_\eta$ | 4.359 | 1.698 | 2.179 | 3.976 | 8.680 |
| $A_f$ | -0.362 | 0.053 | -0.463 | -0.364 | -0.252 |
| $A_\eta$ | -0.530 | 0.627 | -1.531 | -0.617 | 0.860 |
| $I_f$ | 0.569 | 0.122 | 0.380 | 0.548 | 0.855 |

**Table 3:** Median of estimated activation energies of maximum feeding rate (equation 7) for the ciliate predator *Tetrahymena pyriformis* adapted to 15°C, 20°C and 25°C for approximately 20 generations.

| adaptation temperature | activation energy |
| --- | --- |
|  | maximum feeding rate |
| 15°C | 0.663 |
| 20°C | 1.054 |
| 25°C | 1.431 |

## Results

We found that over the course of 20 generations, predator body sizes decreased with increasing adaptation temperature, and thus predators adapted to higher temperatures had smaller average body sizes than predators kept at lower temperatures (Figure 3a). The half-saturation density (Figure 3b) generally increased with experimental temperature with no significant differences for predators adapted to different temperatures and a high variability (Table 2). According to the WAIC, model 3 (equations 6a, 5b) represented our data best, therefore, there was no interactive effect of experimental and adaptation temperature on half-saturation density. The effect of experimental temperature on half-saturation density equaled the activation energy *E*_*ƞ*_ = 4.359 (with a standard deviation of 1.698 and a CI from 2.179 to 8.680) for predators adapted to all three adaptation temperatures. The effect of adaptation temperature on half-saturation density was slightly negative, but insignificant (Table 2). As half-saturation densities should not be affected by body size (Hansen *et al*., 1997) the direct effect of adaptation equaled the realised effect (see Figure 2). Attack rates decreased with experimental temperature, with attack rates of cold adapted predators decreasing faster than attack rates of warm adapted temperature (Supporting Information Figure 9 and Table 4, 5).

In order to calculate the direct effect of adaptation on maximum feeding rates (Figure 2d), we factored body size into the respective maximum feeding rates, a posteriori to the per capita estimation of functional response parameters (equation 8 and 9). Investigating the direct effect of adaptation on maximum feeding rates revealed that warm adapted predators had highest maximum feeding rates at the highest experimental temperatures and lowest maximum feeding rates at the lowest experimental temperature (Figure 3c). Recent studies suggest a power law scaling of body size close to one for chemo-heterotrophic unicellular organisms yielding the same general results (Okie *et al*., 2016; results shown in Supporting Information, Figure 10). The physiological temperature adaptation was affecting activation energies. In predators adapted to 15°C, the activation energy for maximum feeding rate (equation 6a) was approximately 0.66 and increased with adaptation temperature to approximately 1.05 and 1.43 for predators adapted to 20°C and 25°C, respectively (Table 3). In the realised adaptation scenario, maximum feeding rates (Figure 3d) generally increased with experimental temperature. Over most of the observed range of experimental temperatures, maximum feeding rate was highest for predators adapted to 15°C, followed by those adapted to 20°C, and lowest for those adapted to 25°C. Predators adapted to 25°C showed the steepest increase in maximum feeding rate with increasing experimental temperature, while predators adapted to 20°C and 15°C showed a shallower increase (Figure 3d). This resulted from a positive interaction between experimental and adaptation temperature (Table 1). However, at 25°C experimental temperature, there was no difference between maximum feeding rates of predators adapted to 15°C, 20°C or 25°C.

## Discussion

Increasing temperatures are putting a strain on biodiversity in ecosystems world wide. Previous studies have revealed an increasing mismatch between maximum feeding rates and metabolism with warming as an often overlooked and until recently poorly understood cause of extinction (Vucic-Pestic *et al*., 2011; Rall *et al*., 2012; Fussmann *et al*., 2014).

Here, we investigated the effect of possible temperature adaptations on feeding interactions. After an adaptation period of approximately 20 generations, predator body size had decreased significantly for predators adapted to 25°C compared to predators kept at the lowest adaptation temperature according to our prediction based on previous studies (Bergmann, 1847; Daufresne *et al*., 2009; Yvon-Durocher *et al*., 2011). We ran functional response experiments along a temperature gradient with predators adapted to different temperature regimes and found that experimental temperature has an effect on half-saturation densities of predators adapted to all three adaptation temperatures (Fussmann *et al*., 2014). Using more than one predator per experimental treatment, the particularly high values of half-saturation densities might be explained by predator interference. Interference has been observed among unicellular organisms (Curds & Cockburn, 1968) and can be affected by temperature changes (Lang *et al*., 2012). By reducing the time available for prey encounters, interference lowers the feeding efficiency of predators (Abrams & Ginzburg, 2000). In cases where half-saturation density and interference both increase with warming, this could lead to a combined effect on half-saturation densities. Declining attack rates with experimental temperature can be caused by increasing interference and therefore corroborate this assumption. However, since we did not vary predator density to manipulate the strength of predator interference, this can only be speculated. According to our model comparison there is no interactive effect of experimental and adaptation temperature on half-saturation density. This suggests, that the effect of adaptation temperature on half-saturation-density is buffered by a simultaneous temperature adaptation of attack rate and maximum feeding rate. Therefore, adaptation of half-saturation densities should be excluded as a possible mechanism to counteract temperature effects on carrying capacities and decreasing prey abundances at higher temperatures in natural systems. However, predators adapted to higher temperatures show the steepest increase of maximum feeding rate with increasing experimental temperature enabling them to react to increasing temperatures quicker and increase their energy intake faster within the measured temperature range. Predators adapted to 25°C show lower maximum feeding rates at 15°C and 20°C than cold adapted predators, while at 25°C experimental temperature, all predators show similar maximum feeding rates. In our experiment we were unable to document potential changes in metabolism for predators adapted to different adaptation temperatures, which leaves two possible hypotheses to explain our findings. The hypothesis that metabolic rates were unaffected by adaptation temperature leads to the conclusion that predators adapted to higher temperatures have gained a disadvantage at lower experimental temperatures becoming less efficient compared to their cold adapted counterparts, while there is no clear advantage gained at high experimental temperatures. However, due to smaller body sizes of warm adapted predators, predators adapted to 25°C adaptation temperature are expected to have lower metabolic demands compared to cold adapted predators, if any potential physiological adaptation of metabolism is taken into account (Brown *et al*., 2004). Relevantly, our experimental units contained more than one predator individual, these lowered metabolic demands can lead to an increase in predator interference, reducing maximum feeding rates at low experimental temperatures. With increasing experimental temperatures, these predators will prioritise feeding over predator interaction leading to the strong increase of maximum feeding rates with experimental temperature in warm adapted predators. While some studies predict activation energies for maximum feeding rates ranging from 0.6-0.7 eV, our results are in the range of activation energies reported for ciliated protozoan and other unicellular organisms around 0.772 eV (Hansen *et al*., 1997; Vasseur & McCann, 2005). Activation energies for metabolism drawn from respiration measurements by Laybourn & Finlay (1976) of 0.96 eV (Fussmann *et al*., 2014), match the range of activation energies of maximum feeding rates in our functional response measurement. Predators adapted to 20°C and 25°C show higher activation energies to counteract increasing metabolic demands at higher temperatures. Combining the strong increase in maximum feeding rate with the change in intercept caused by body size adaptation, these results are in line with the hypothesised interactive effect of body size adaptation, and adaptation temperature and experimental warming on maximum feeding rates.

Over the timespan of 20 generations, our results as well as previous studies have shown that adaptation to increased temperatures influences protist body sizes (Atkinson *et al*., 2003) highlighting the importance of trans-generational studies regarding not only genetic adaptation but also phenotypic changes (DeLong *et al*., 2016). Larger species, predominantly found at higher trophic levels (Riede *et al*., 2011) are most vulnerable to extinction due to an energetic mismatch with increasing temperatures (Binzer *et al*., 2012). This leads to a shift towards smaller species in aquatic systems (Daufresne *et al*., 2009; Yvon-Durocher *et al*., 2011). The relationship between increasing predator body size and maximum feeding rate follows a ¾ power-law scaling (Hansen *et al*., 1997; Rall *et al*., 2012), leading to lower maximum feeding rates in smaller predators. To disentangle the indirect effect of predator body size on the realised maximum feeding rate from the direct effect of physiological adaptation we corrected our results accordingly (see figure 2 and equations 8 and 9 for a detailed derivation). Once this change in body size is accounted for, we found that at 25°C experimental temperature, maximum feeding rates shift towards a scenario that suggest a specialised temperature adaptation of predators. While at 15°C predators adapted to that temperature show the highest maximum feeding rates, at 25°C predators adapted to 25°C show the highest maximum feeding rates. There is not one culture adapted to have the best fitness at the full temperature range, rather predators seemed to be adapted to their respective temperature. The direct physiological adaptation of maximum feeding rates leads to a stronger increase in maximum feeding rate with experimental temperature in warm adapted predators. Further, in form of a morphological adaptation to warming, with a smaller average body size, predators increase per-biomass consumption while reducing metabolic demand. This increases the effect of physiological adaptation of maximum feeding rates, resulting in a combined effect increasing overall energy efficiency in warm adapted predators at high temperatures.

In conclusion, our results suggest that while un-adapted predators face a mismatch between maximum feeding rates and metabolic demands with increasing temperatures leading to starvation and extinction of predators, adaptation poses a viable escape from this scenario. By decreasing their body size over the course of 20 generations at higher temperatures, predators lower their per-capita metabolic rates. Therefore, the ratio between metabolic costs and maximum feeding rates increases for warm adapted predators, decreasing their risk of starvation. The decrease in the risk of starvation also implies a decreased risk of extinction which may buffer expected biodiversity loss with climate warming and increased ecosystem stability.

It is widely accepted that adaptation occurs within ecological time spans and is therefore of utmost importance for the understanding of population stability and ecosystem dynamics under the threat of an increasingly fast changing environment (Holt, 1990; Lynch & Lande, 1993; Burger & Lynch, 1995; Merilä, 2012; Merilä & Hendry, 2014). Especially in a homogeneous environment like water, where stressors cannot be avoided by migration or refuge in microhabitats, the strain of climate change poses a particularly high risk for populations (Bergmann *et al*., 2010). Adaptation might be a possible way for populations to deal with increasing temperatures and persist in a warming environment.

## Acknowledgements

K.E.F. received funding from the Dorothea Schlözer Programme of Göttingen University. K.E.F., B.R., U.B. and B.C.R. gratefully acknowledge the support of the German Centre for Integrative Biodiversity Research (iDiv) Halle-Jena-Leipzig funded by the German Research Foundation (FZT 118). We would like to thank A.Binzer and D.Perkins for helpful suggestions and comments.

